# Non-Invasive Quantification of Viability in Spheroids Using Deep Learning

**DOI:** 10.1101/2025.03.09.642246

**Authors:** Daniel Dubinsky, Shahar Harel, Amir Bein, Abraham Nyska, Sarah Ya’ari, Baran Koc, Faiza Anas, Isaac Bentwich, Lior Wolf

**Author notes:** Corresponding Author: Daniel Dubinsky.

## Abstract

In vitro viability assays are widely used in drug discovery, development, and pharmacovigilance. Traditional methods for evaluating cell viability often involve destructive processes, rendering the culture non-viable. As such, these methods are suitable only as endpoint solutions, providing a single measurement per cell culture and precluding further analyses of the cells. In this study, we introduce Neural Viability Regression (NViR), a real-time, deep learning-based method that enables non-invasive quantification of culture viability using microscopy images. The non-intrusive nature of NViR allows for frequent viability evaluations throughout experiments, capturing subtle changes while maintaining the structural integrity of the culture and significantly reducing both culture and labor costs. We demonstrate NViR’s applicability by using it to predict Drug-Induced Liver Injury (DILI) in known drugs. By exposing human liver spheroids to 108 FDA-approved drugs and capturing microscopy images over time, NViR’s viability assessments accurately predict whether a drug induces DILI in humans, playing a critical role in enhancing liver safety protocols. The cost-effectiveness and non-invasive characteristics of NViR enable high-frequency, high-throughput viability assessments. Consequently, NViR is poised to reduce both the costs and incidences of failures in drug discovery and development.

## 2 Main

Adenosine triphosphate (ATP) is a molecule that serves as the primary energy source in cells [1, 2]. It plays a crucial role in various biological processes, including cellular metabolism, muscle contraction, and signal transduction [1]. Apart from its role in cellular energy transfer, ATP has gained significant attention as a biomarker for many applications [3, 4, 5, 6, 7, 8, 9].

One notable application is the assessment of cell viability, since ATP levels indicate the overall state of a cell’s metabolic and mitochondrial activity [2, 1]. Viable cells maintain high ATP levels, while a decline in ATP is often associated with viability degradation, cellular damage, apoptosis, necrosis, or metabolic dysfunction [10]. Viability assays are widely used for drug discovery and development and thus are significant tool in laboratories worldwide [11]. For example, viability can be used as the main indicator for drug-induced liver injury (DILI) [8, 12]. Other diagnostic settings include assessing the hepatotoxic potential of non-pharmaceutical chemicals [13], conducting tumor chemosensitivity testing [14], estimating mitochondrial toxicity [6, 9], and identifying the cytotoxicity of anticancer drugs [7].

The bioluminescence method is frequently used to assess viability, relying on the quantification of intracellular ATP levels, and is widely recognized as the gold standard [15, 7, 16, 17, 18, 19]. It is an accurate and consistent method [18] for evaluating the total ATP content within a given sample.

While the bioluminescence viability quantification assay is widely accepted and used due to its numerous advantages, its main, critical drawback is the requirement of performing a lysis process that irrevocably terminates the viability of the target cells or tissues [5]. As such, the bioluminescent assay is an endpoint assay with a single application per spheroid. This limitation prevents observing subtle changes over time using the same spheroid, thereby potentially hiding insights. To partially overcome this limitation, researchers generate multiple biological repetitions, by repeating the viability assay on multiple spheroids under the same conditions [14, 13]. While this approach partially addresses the issue, generating nearly-identical repetitions is both biologically challenging [20, 21] and expensive, requiring additional scientists and spheroids. In practice, because of these limitations, researchers typically run a low number of viability quantification endpoints per experiment (usually 1-3) [21, 8, 7].

Real-time viability assays, on the other hand, enable continuous monitoring of cell health without requiring lysis. A prominent example is the RealTime-Glo^TM^ MT Cell Viability Assay [22], which employs an engineered luciferase and a pro-substrate added directly to the culture medium. In this assay, metabolically active cells convert the pro-substrate into a substrate for luciferase, resulting in a measurable luminescent signal. However, a limitation of this method is the depletion of the prosubstrate by metabolically active cells, which affects the duration for which the luminescent signal correlates linearly with cell numbers [23]. Additionally, environmental and chemical factors can impact the assay’s results, although the reagents’ effects are typically minimal, if present at all. Despite these limitations, RealTime-Glo remains a widely utilized assay in the industry [24, 25, 26]. In contrast, our method, based solely on bright-field imaging, not only retains the continuous monitoring capabilities of real-time assays, but also significantly advances them. NViR overcomes key limitations such as substrate depletion and environmental susceptibility, offering a more robust and reliable approach to viability quantification. Moreover, NViR’s ability to analyze images for subtle viability changes without invasive procedures presents a groundbreaking improvement in both methodology and data integrity. This represents a considerable leap forward in cell viability assessment,

Spheroids exhibit morphological alterations when damaged[27]. Figure 1 shows bright-field images of spheroids with their corresponding viability levels on top. Notably, lower viability spheroids present distinct features ranging from altered contours and varied nucleus darkness to complete breakdown. Given these observations, we hypothesize that both visible and subtle features can be detected by deep learning algorithms and used to predict viability from images.

**Figure 1:**
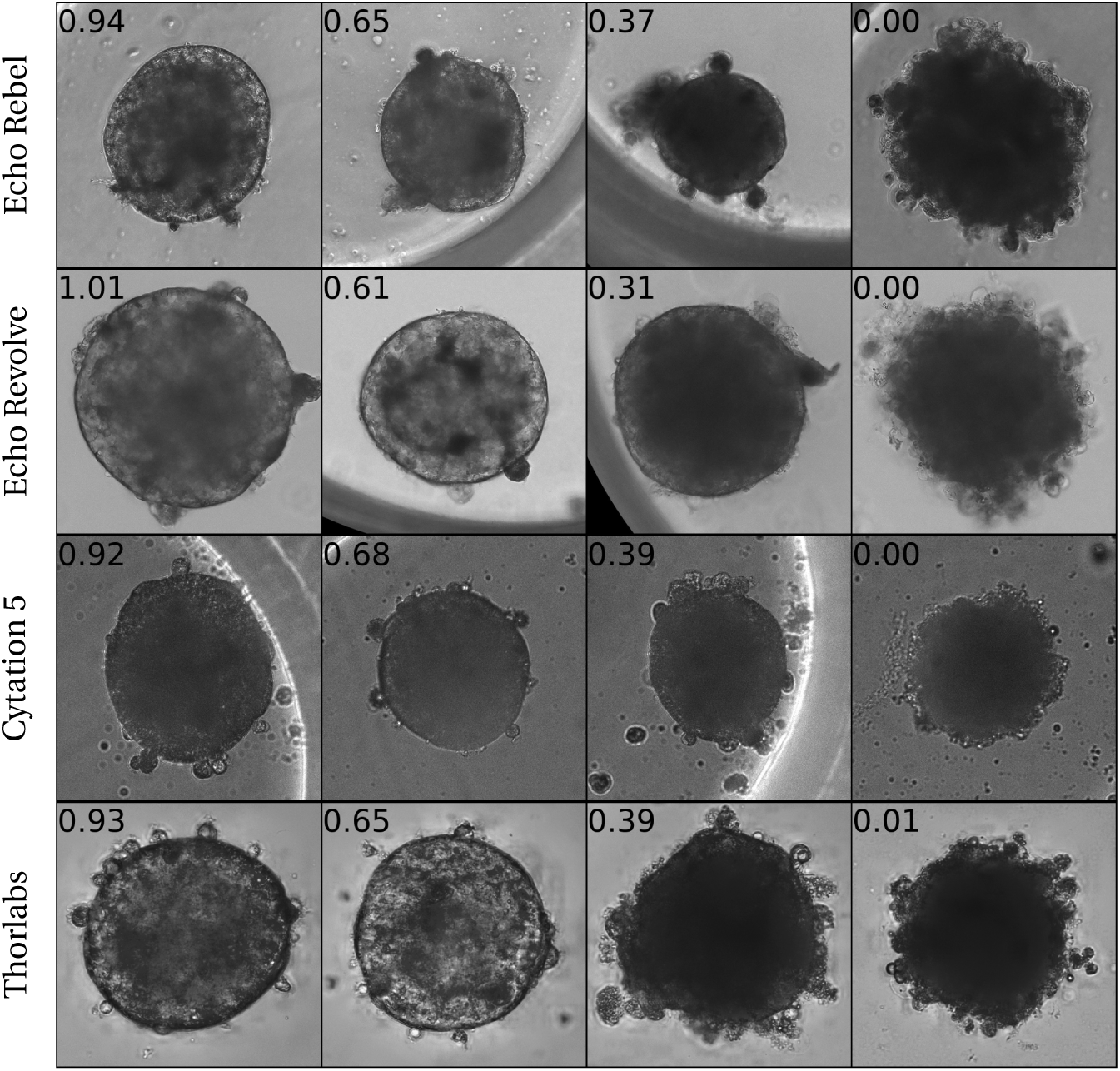
Bright-field images of 16 liver spheroids captured by 4 different microscopes with varying viability along with the ATP(measured using CTG) values normalized by respective control. The loss in viability is associated with a range of morphological abnormalities, ranging from altered contour shapes and varied nucleus darkness to complete breakdown.

In this work, we present a non-invasive deep learning method for assessing viability, which addresses the primary limitation of bioluminescence assays and exhibits high predictive accuracy. Our approach includes a data generation process that yields a labeled training dataset of spheroids, and a deep learning technique for estimating viability levels from the spheroid images. Figure 2 offers an overview of our approach; the upper section depicts the data generation process, and the lower section demonstrates the inference process using a deep learning regressor.

**Figure 2:**
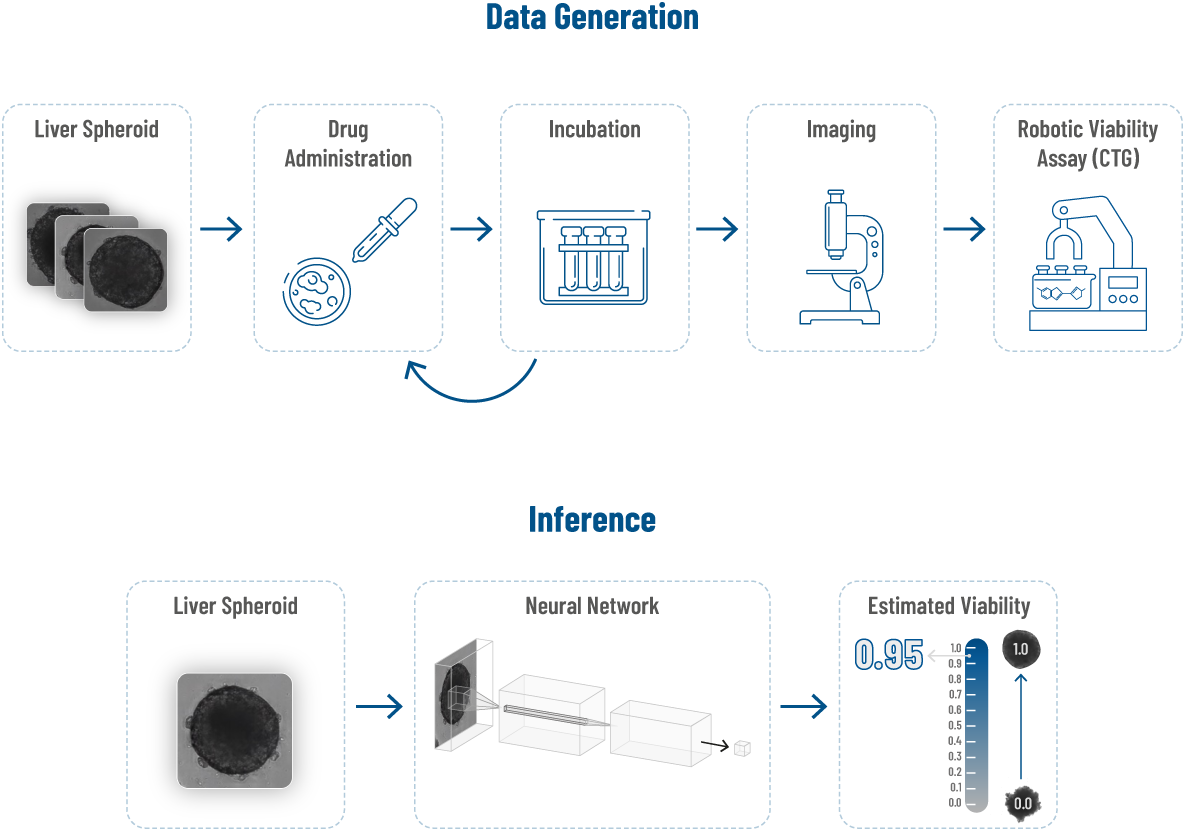
Overview of the proposed framework. The top part illustrates the data generation process, and the bottom part illustrates viability prediction based on a spheroid image.

We used a high-throughput microphysiological system (MPS) platform to generate our dataset. The MPS platform manages spheroid treatments, oversees the experiment, and conducts the ATP quantification assay.

From this process, we generated a dataset containing 17,704 unlabeled and 2,359 labeled spheroid bright-field images, each paired with its measured viability level. We have made this dataset publicly available to support further research in the community.

The neural network we employ on top of this dataset is based on well-established ResNet architecture [28] followed by a Multi-Layer Perceptron (MLP) as a regression head. The training process itself is also a typical one.

The resulting model exhibits strong generalization capabilities in both in-domain and out-of-domain settings, effectively overcoming lab-driven deviations from the original distribution. In particular, our model achieves consistently high performance, even when provided with images captured using a microscope different from the one used to generate the training data. This adaptability promotes the method’s wide applicability across a range of diverse experimental conditions.

Overall, NViR enables non-invasive, high-throughput estimation of viability based solely on image data, saving both spheroids and time while providing more detailed insights. Our experiments demonstrate that NViR outperforms both computational baselines and human experts, specifically pathologists, who are typically employed for similar tasks [29, 30, 31]. We delve into the latent space of NViR, an intermediate representation into which NViR encodes the most meaningful visual features, exploring the morphological semantics of this representation. Finally, we showcase a practical real-world application by using NViR to predict liver toxicity.

## 3 Results

In this section, we present the comprehensive results of our study. The section is structured into three key subsections, the first of which focuses on the rigorous comparison of NViR with several baseline approaches. The second subsection delves into the interpretability of NViR, revealing how it identifies factors that contribute to viability estimations. The final subsection illustrates NViR’s practical application in drug-induced liver injury (DILI) prediction, underscoring its potential in advancing non-invasive cell culture analysis and drug safety assessment.

### 3.1 Viability Quantification

We evaluate the performance of NViR by comparing it with human experts and algorithmic baselines. To ensure a robust evaluation, considering the natural variations in data due to differences in equipment, staff variances, and biological factors within laboratory operations, we conduct evaluations across four distinct domains: *A*, *B*,*C*, *D*. These domains are distinguished by either physical location (i.e., different laboratories) or the type of microscope used. For training, we use a single domain, denoted as *A*, and implement a time-based train/test split, reserving the most recent studies for evaluation.

As an algorithmic baseline, we introduce the Classical Vision Encoder (CVE), which combines computer vision algorithms to analyze visual characteristics from both the spheroid image and its segmentation mask. These characteristics include metrics such as the ratio of circumference to perimeter, curvature, and relative brightness in relation to the background. Subsequently, a machine learning regressor is trained to estimate viability levels based on these extracted features.

An additional baseline is established by utilizing the assessment of a pathology expert. Given that pathologists often evaluate the viability and damage of cell cultures [29, 30, 31], their insights serve as a meaningful baseline. The pathologist employed a morphology-based rating system to score spheroid images according to their viability. Scoring was based on three properties linked to viability: cytoplasmic changes [32], nuclear changes [33], and spheroid contour shape changes [34, 19]. A spheroid was given three scores in the range 0*−−*4, one for each property. All pathologist scores are relative to the control spheroid. A machine learning-based regressor is then trained on these scores to predict viability levels. More details about the pathologist’s labeling process can be found in the methods section.

A third baseline is obtained by combining the classical computer vision features provided by the CVE method, and those provided by the human pathologist. A machine learning-based regressor is trained on the concatenation of these features to predict viability levels.

Tables 1 and 2 provide a comprehensive breakdown of the R-squared and mean absolute error values (MAE) for all methods employed across the domains. Figure 3 depicts scatter plots comparing the predicted and actual values.

**Table 1:** Mean and Standard Deviation for *R*^2^ (higher is better) of NViR and the baselines across four domains, with all models trained exclusively on domain *A*.

| Model | Domain |  |  |  |
| --- | --- | --- | --- | --- |
|  | A | B | C | D |
| CVE (baseline) | 0.16 ± 0.00 | 0.56 ± 0.00 | 0.32 ± 0.00 | 0.68 ± 0.00 |
| Pathology (baseline) | 0.53 ± 0.03 | 0.69 ± 0.00 | 0.39 ± 0.03 | 0.59 ± 0.03 |
| Pathology + CVE (baseline) | 0.56 ± 0.00 | 0.70 ± 0.00 | 0.50 ± 0.00 | 0.62 ± 0.00 |
| NViR | **0.83 ± 0.03** | **0.84 ± 0.06** | **0.70 ± 0.06** | **0.80 ± 0.05** |

**Table 2:** Mean and standard deviation of mean-absolute-error (MAE; lower is better) for NViR and the base-lines across four domains, with all models trained exclusively on domain *A*.

| Model | Domain |  |  |  |
| --- | --- | --- | --- | --- |
|  | A | B | C | D |
| CVE (baseline) | 0.253 ± 0.000 | 0.215 ± 0.000 | 0.230 ± 0.000 | 0.166 ± 0.000 |
| Pathology (baseline) | 0.209 ± 0.006 | 0.185 ± 0.000 | 0.219 ± 0.006 | 0.193 ± 0.007 |
| Pathology + CVE (baseline) | 0.194 ± 0.000 | 0.181 ± 0.000 | 0.195 ± 0.000 | 0.200 ± 0.000 |
| NViR | **0.113 ± 0.012** | **0.134 ± 0.033** | **0.151 ± 0.018** | **0.135 ± 0.024** |

**Figure 3:**
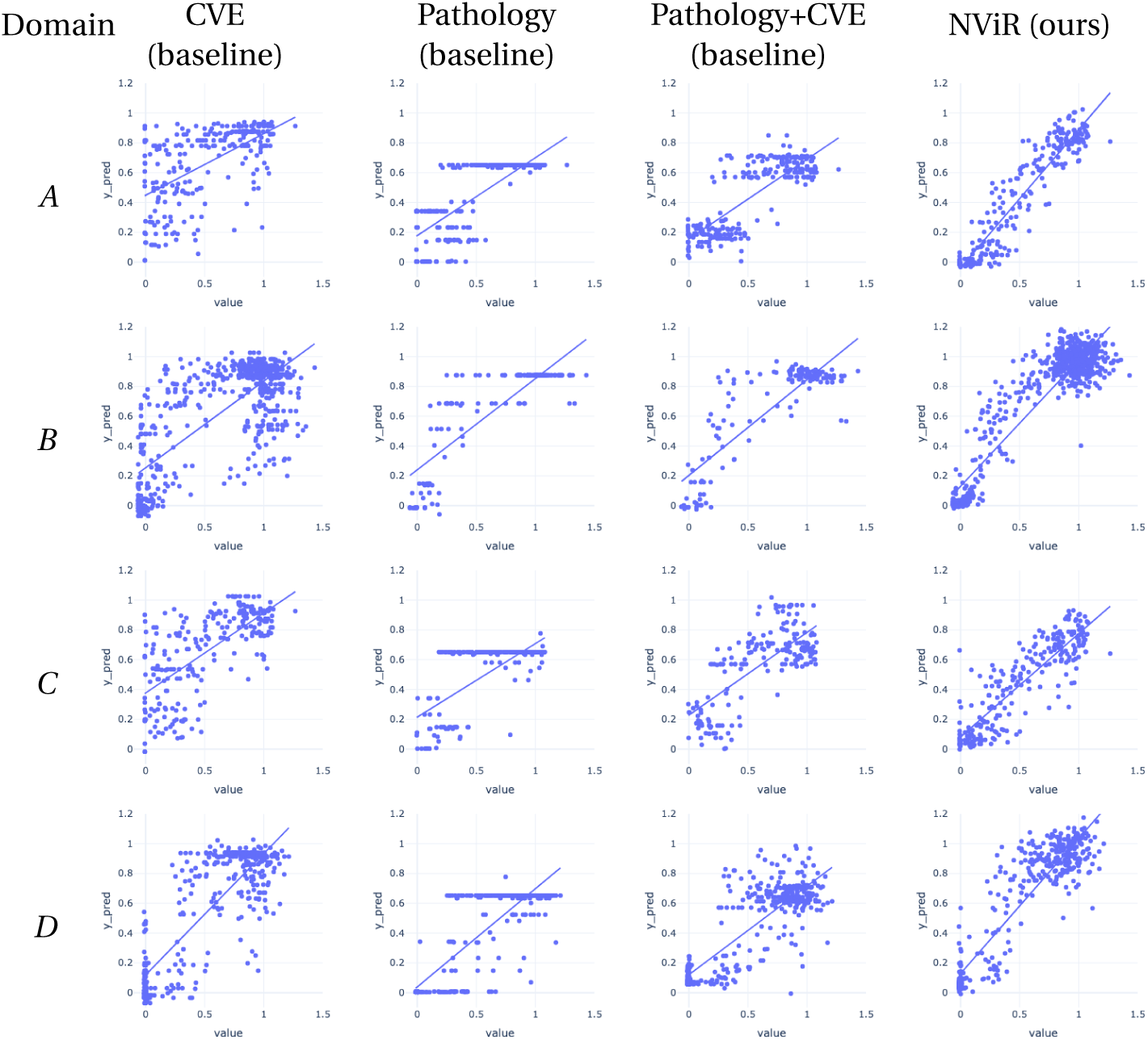
Comparison of predicted viability (y-axis) against ground truth viability (x-axis) for different methods and domains. Each row represents a domain (A, B, C, D), and each column corresponds to a different method (CVE baseline, Pathology baseline, Pathology+CVE baseline, and NViR). The scatter plots for each method within a domain visually illustrate the accuracy of viability predictions compared to the actual values.

Among the four approaches, the CVE method exhibits the lowest performance. The CVE feature space consists of manually crafted, domain-specific features, in contrast to the end-to-end feature learning of the deep learning method.

Next, the pathology baseline, while achieving non-trivial performance and being superior to the CVE baseline, still leaves room for improvement. The pathologist ranking system is limited to only 125 potential combinations, spanning the three parameters, with five levels each. Notably, 64% of the labels correspond to just three of these combinations. Such a limited feature space poses a significant challenge for estimating continuous viability. In Figure 4, we present the three main combinations of parameters employed by the pathologists, each associated with a wide spectrum of continuous viability values.

**Figure 4:**
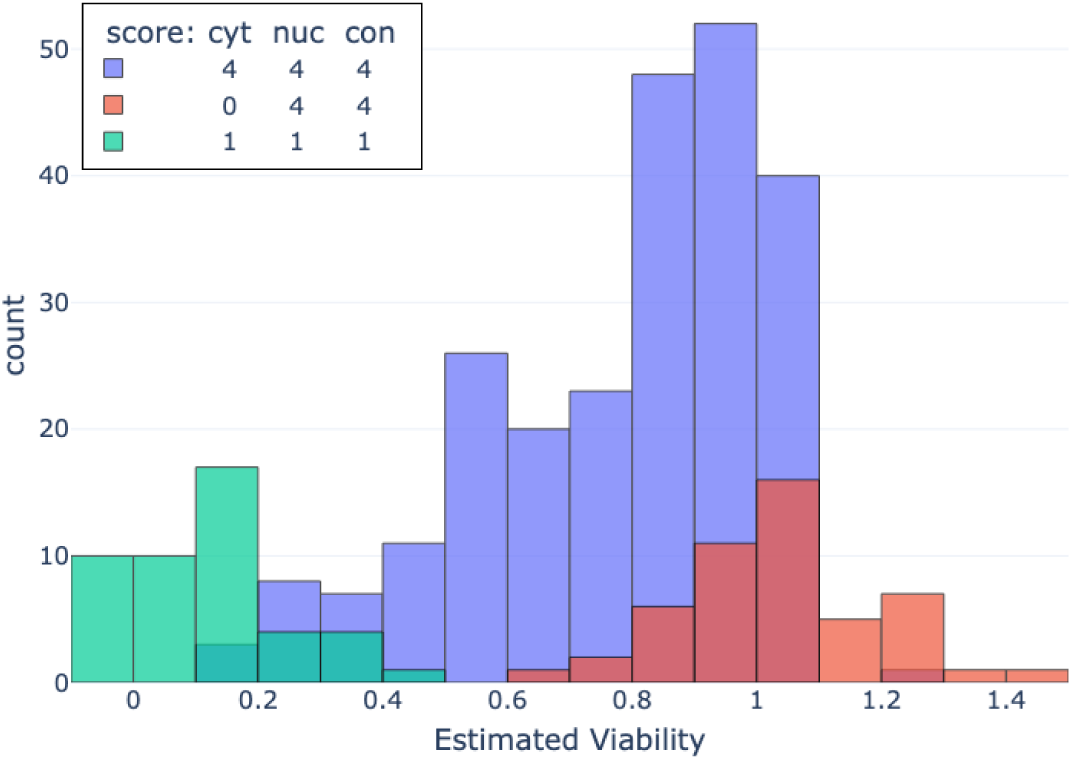
Control-Normalized ATP values of spheroids that received the three most frequent pathology score combinations. Each combination comprises cytoplasm changes score, nuclear changes score, and contour changes score.

The combination of both pathologist annotations and the CVE yields notably improved results compared to their individual performance. This confirms our initial hypothesis that image data contains valuable information related to viability. Furthermore, the increase in performance indicates that computational features can provide additional value on top of the pathologist method.

NViR exhibits marked improvement over all these baseline approaches, achieving significantly better results. Its ability to generalize to domains not seen during training underscores the potential of NViR as a precise and reliable method for quantifying viability.

### 3.2 Interpretability

We aim to interpret what NViR encodes for predicting viability by identifying connections between NViR’s latent space, which encodes images, and established biomarkers with semantic significance. NViR’s latent space is a 512-dimensional vector space into which it encodes images. A high correlation between a coordinate in the latent space and a biomarker indicates that NViR captures information similar to the biomarker, advancing our understanding of NViR.

For biomarker analysis, we utilize fluorescence staining — a technique commonly used in toxicity assessments to detect and monitor cellular damage. We employed ChromaLive staining[35], a multi-chromatic dye with two excitation and three emission wavelengths, which is responsive to phenotypic changes such as endoplasmic reticulum stress, apoptosis, and autophagy[35]. After acquiring pairs of brightfield and stained images for 71 spheroids, three of which are shown in Figure 5, we calculated the biomarker score as the sum of intensities normalized by the spheroid area in the stained image. Additionally, we derived NViR feature vectors by encoding the bright-field images into the latent space.

**Figure 5:**
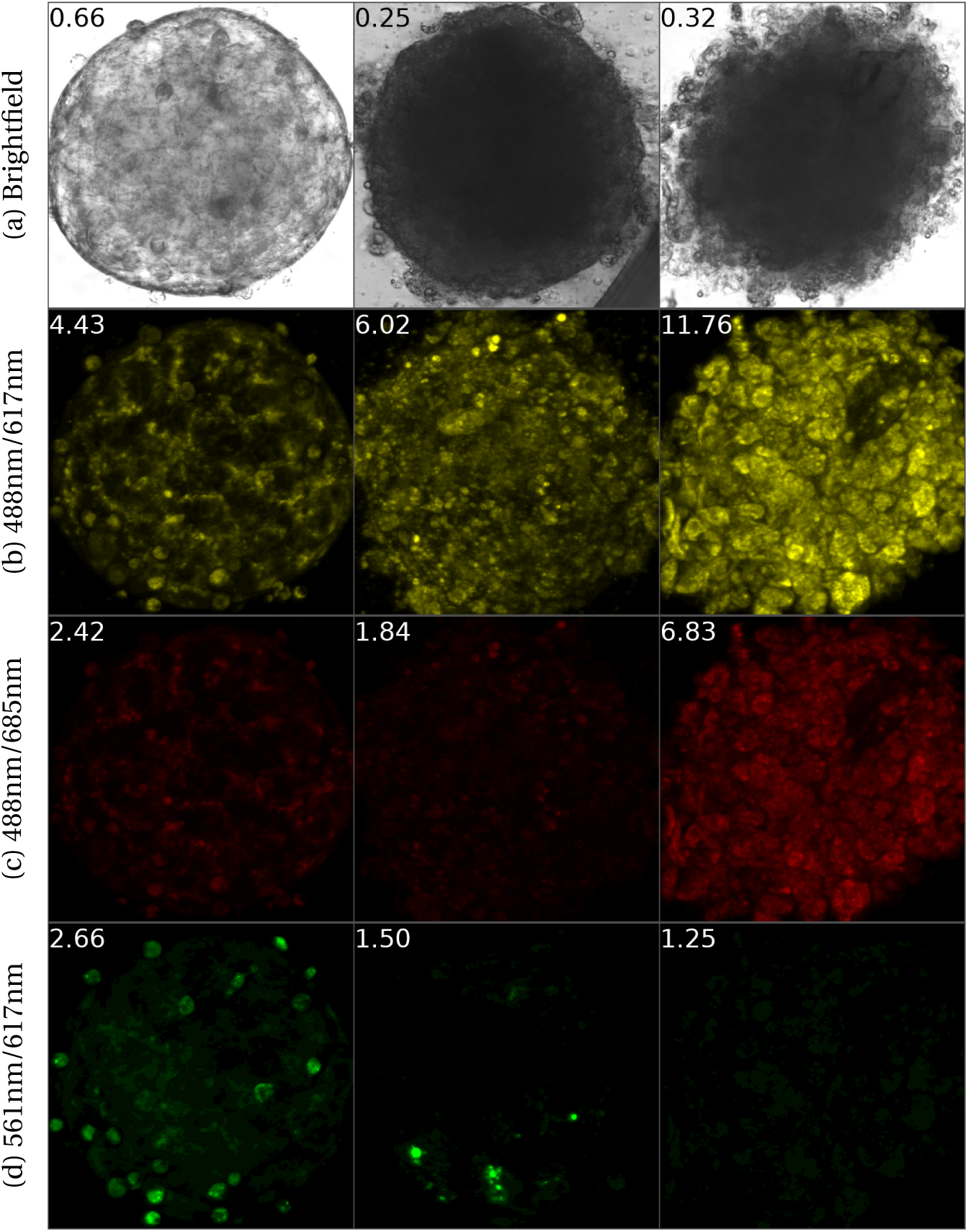
Three Chromalive-stained spheroids of varying viability, where each column represents a spheroid. The row (a) shows bright-field images taken with the Thorlabs microscope, while rows (b-d) are fluorescence images taken with the Yokogawa CQ1 confocal microscope at three excitation/emission lengths: 488nm/617nm (yellow), 488nm/685nm (red), and 561nm/617nm (green). The numbers at the top left of each image indicate the viability for row (a) and the calculated biomarker for rows (b-d).

Upon computing the biomarker values from the stained images, we assessed the absolute Pearson correlation between each biomarker and the latent space of the model. Similarly to most neural models, NViR is not optimized to create a disentangled latent space. Therefore, Principal Component Analysis (PCA) is used to extract uncorrelated directions in the latent space, sorted by variance. Specifically, we examine the top 10 principal components (PCs) of NViR’s latent space vectors. For the biomarkers, we consider the sum of the intensities of the stained image for each image and each staining method.

The first PC, accounting for 42% of the variance in NViR’s latent space, shows a high correlation to both viability (*cor r =* 0.86; *P <* .001) and the biomarkers (*cor r >* 0.61; *P <* .001). As it turns out, the three staining biomarkers are also correlated with viability (in all cases *corr >* 0.76; *P <* .001). To explore beyond this connection, we examine the subspace of the 10 PCs that are mostly orthogonal to viability. Specifically, we consider the 8 PCs with less than 0.1 correlation to viability.

Out of the 8 PCs, some exhibit non-trivial correlations to the biomarkers:

- **Chromalive 488nm/617nm(yellow)** - correlated with PC8 0.67, *P <* .001 (p-values are computed using a permutation test with 1000 repeats, considering the highest correlation with any of the eight PCs, the p-values of the Pearson correlation is always before 10*^−^*^4^.)
- **Chromalive 488nm/685nm(red)** - correlated with PC5 0.56, *P <* .001
- **Chromalive 561nm/617nm(green)** - correlated with PC8 0.66, *P <* .001

These results suggest that NViR, while trained to predict a specific ATP-based vi-ability, is also highly correlated with the information obtained by the staining techniques.

### 3.3 Liver Toxicity Prediction

Assessing the safety of drug candidates is a fundamental aspect of the drug development process [36], serving as a critical protective measure for both patients and the pharmaceutical industry. Among the primary safety concerns associated with drugs is the risk of liver toxicity, specifically known as Drug-Induced Liver Injury (DILI), which is a major cause of drug withdrawal and failure in clinical trials [37, 38]. To predict potential DILI, cell culture viability assessment is widely employed in drug safety evaluations [39, 12, 29, 8].

Proctor et al. [8] introduce a method for DILI prediction using the bioluminescent CellTiter-Glo(CTG) [5] assay for viability quantification. To validate their method, a subset of 110 drugs from Garside et al. [40] is used, labeled according to their binary DILI label.

To determine the DILI label for a given drug, they expose liver spheroids to varying concentrations of the drug, and viability data are collected after a 14-day incubation period. Subsequently, the researchers calculate the drug’s half maximal inhibitory concentration (*IC*_50_), which represents the drug concentration causing a 50% reduction in viability levels compared to control levels.

In addition to *IC*_50_, another parameter employed is *C_max_*, which denotes the maximum concentration of the drug in the plasma following administration. A DILI label is then determined using a predefined threshold, considering either the *IC*_50_ value or the Margin of Safety (*MOS*), which is defined as 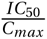.

We demonstrate that NViR is capable of replacing the endpoint bioluminescent assay used by [8] and can thus be used for identifying liver toxicity, while eliminating the need for the bioluminescent assay and the sacrifice of spheroids, relying solely on microscopy images. Moreover, conducting the assay in a non-invasive manner enables us to carry it out multiple times within a single experiment, providing a more detailed perspective on viability over time.

We scanned 108 out of the 110 drugs at various concentrations for a 7-day period, capturing daily bright-field images of the spheroids. The scans took place at sites A and B, the imaging was performed using the Thorlabs microscope, and information on the drugs used in this study are provided in Supplementary Table S1. Using NViR, we assess the viability levels from these images each day and subsequently calculate the daily *IC*_50_ values of the drugs. *IC*_50_ is then divided by *C_max_* to create the approximate NViR-based *MOS* value per day.

Following Proctor et al., we use cutoffs over NViR-based *MOS* values for DILI classification. Figure 6 compares the ROC curves obtained by using thresholds on day 7 *MOS* values for DILI classification (we did not continue the experiment past day 7). Evidently, NViR’s performance on day 7 is on par with the approach presented by Proctor et al. after 14 days of incubation. This indicates that NViR’s ability to closely approximate bioluminescence-based viability measurements translates effectively to liver toxicity prediction.

**Figure 6:**
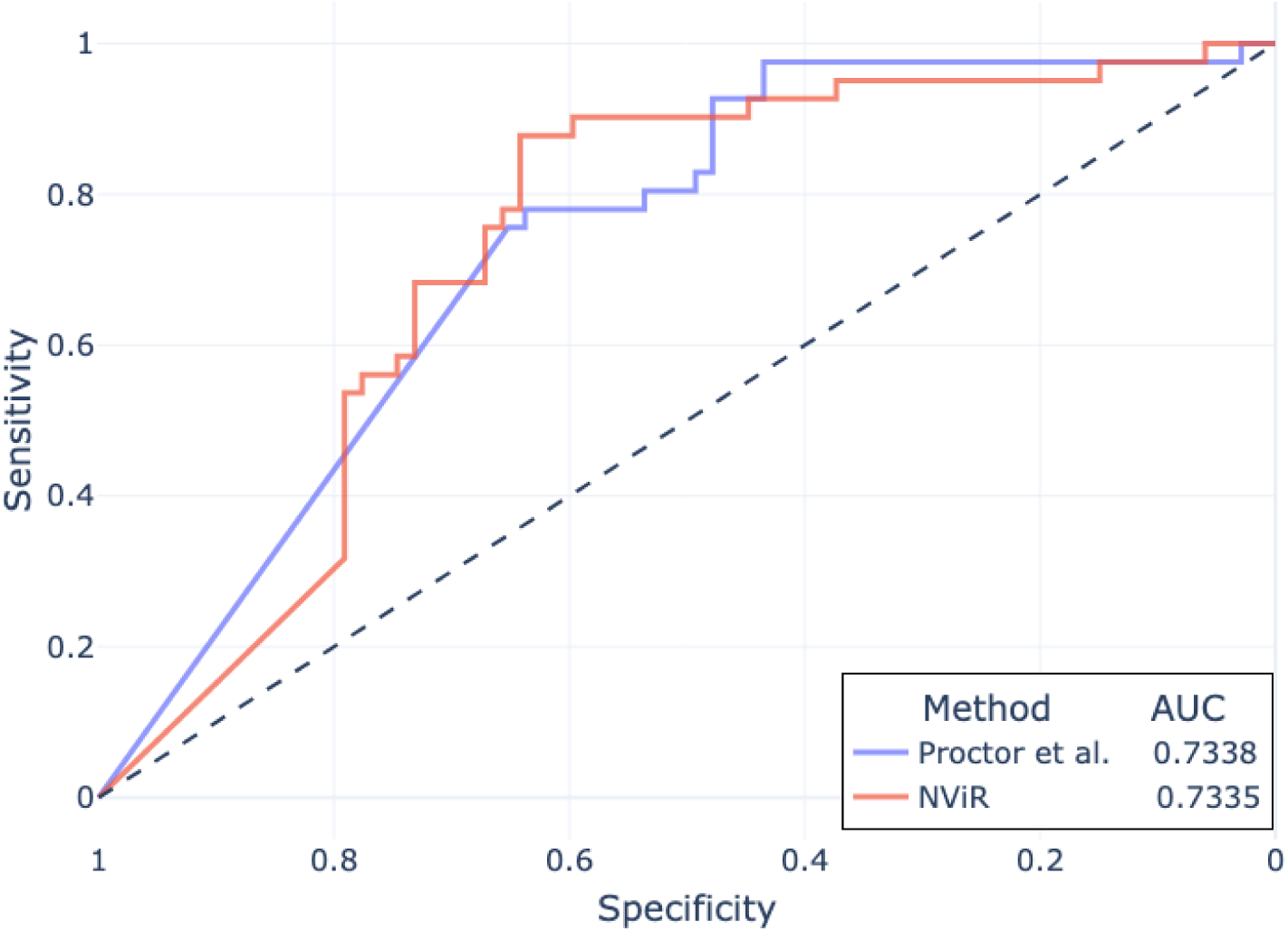
ROC curves of DILI classification with *MOS* values calculated using NViR and CTG by Proctor et al.[8]. The AUC is indicated in the legend.

To provide more insight, we present in Figure 7 two spheroids over the course of the experiment: the spheroid on top was treated with Salbutamol (Albuterol), a drug clinically known to be non-liver-toxic [41], while the drug used on the bottom spheroid, Nefazodone, is known to be highly liver toxic [42]. Both are displayed with normalized viability levels on top. We observe that the non-toxic treatment induces almost no visual changes to the spheroid over time, while the spheroid treated with DILI-positive drug, in a toxic dose, undergoes significant morphological changes.

**Figure 7:**
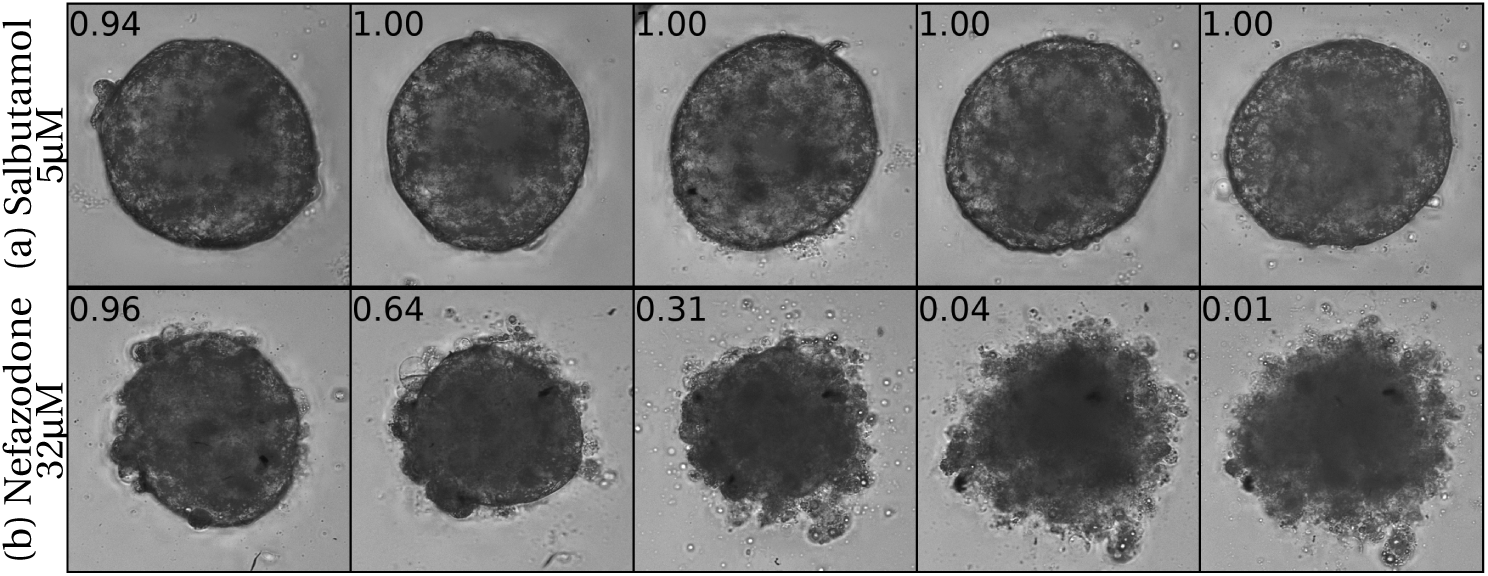
Two spheroids exposed to drugs, shown over the course of the experiment from days 3 to 7, arranged from left to right. (a): Spheroid treated with a non-livertoxic drug(Saburamol). (b): Spheroid treated with a liver-toxic drug(Nefazodone).

Next, we discuss the potential value of observing continuous viability using NViR as opposed to endpoint assays in the context of DILI prediction. Figure 8 provides an example of the insights unveiled through the use of NViR. We showcase the viability of spheroids subjected to various drug treatments over time. Although the viability measured on day 7 appears similar across all treatment groups, viability trends vary significantly. This valuable information would remain concealed if one were to rely solely on an endpoint assay.

**Figure 8:**
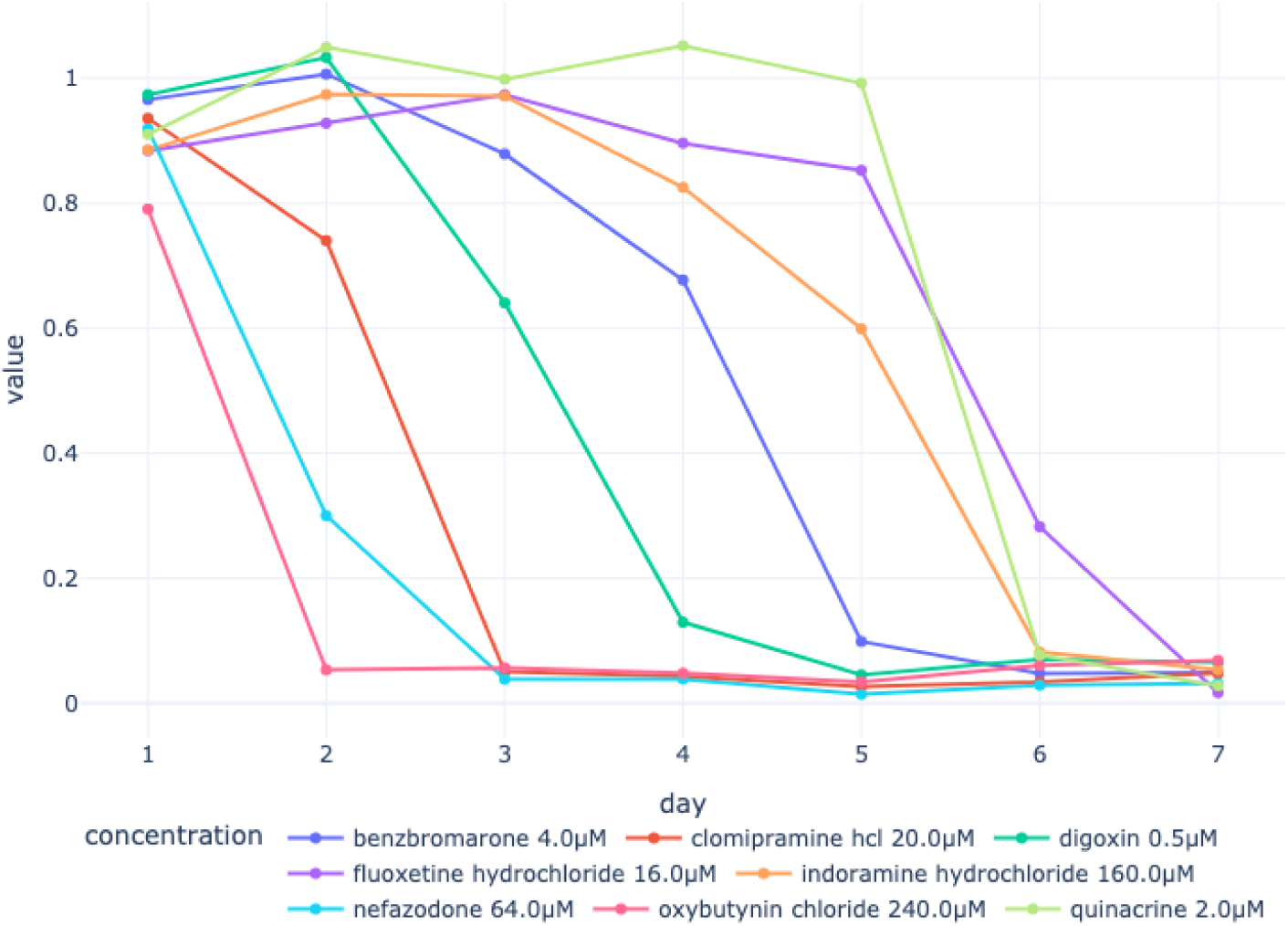
Median daily viability of spheroids subjected to various drug treatments.

## 4 Discussion

In this study we introduced Neural Viability Regression (NViR), a novel, non-invasive, deep learning-based method for assessing the viability of spheroid cultures using bright-field microscopy images. NViR offers significant advantages over traditional viability assays, particularly in terms of continuous, non-invasive monitoring. This advancement positions NViR as a valuable tool in drug discovery and pharmaceutical research, offering a more efficient approach compared to endpoint assays such as bioluminescence.

We demonstrated that NViR’s application as a viability assay extends to clinical scenarios, successfully predicting drug-induced liver injury (DILI) in humans and achieving state-of-the-art performance on a benchmark dataset of known drugs. This underscores NViR’s potential as a valuable tool throughout the drug discovery process, especially for early detection of DILI.

Viability assays extend beyond liver spheroid applications. For example, Omer et al. utilized viability assays on kidney spheroids to assess direct viral-induced cellular injury [43] and to evaluate the dedifferentiation and redifferentiation potential of OCT4-modified kidney cells in renal regenerative medicine [44]. While NViR is trained to predict viability in liver spheroids, it can be adapted for other types of cultures by acquiring similar data to that described in our study and training NViR using the released code. 3D cell cultures are not limited to spheroidal shapes [45]. Since NViR does not rely on specific geometric features, it is likely to be directly applicable to such cultures as well.

The high-throughput and cost-effective characteristics of NViR make it suitable for early integration into the drug development process, particularly in the early phases where thousands of candidates undergo testing [46]. Its non-invasive nature allows for more reliable testing by enabling the use of the same spheroid for multiple measurements obtained at different times. Consequently, NViR has the potential to enhance drug development throughput while simultaneously reducing the occurrence of failures through early detection of drug candidate toxicity.

In future research, we anticipate that the integration of in vitro biology and machine learning will lead to an increased number of AI-enhanced biomarkers. For example, membrane integrity and mitochondrial toxicity are known to have visual, morphological properties [47, 48]. In such cases, deep learning methods have great potential to identify the most important visual properties and potentially correlate them to predict biomarkers based on the image. In our study, we have shown significant correlations between NViR’s latent space and a fluorescence staining technique responsive to specific phenotypic changes. As shown, coordinates in NViR’s latent space can be viewed as AI-enhanced biomarkers.

Beyond vision, other data modalities have been used to create AI-enhanced biomarkers. For example, Azarkhalili et al. condensed mRNA transcription profiles into a low-dimensional latent vector of size 8, which is highly efficient in identifying tissue-of-origin and its cancer type [49]. These biomarkers are expected to reduce time, costs, and human involvement in the research process. They will provide a more comprehensive understanding of changes in cell cultures, increasing the potential to extract additional insights. This is particularly promising in clinical settings, where these biomarkers are expected to serve as early indicators for evaluating drug efficacy and safety.

## 5 Methods

We provide a description of the data collection process, the neural architecture, and the training procedure. All the data, code, and trained parameters can be found in https://github.com/DanielDubinsky/atp_paper

### 5.1 Dataset

#### Data Generation - MPS Platform

A microphysiological system (MPS) is a state-of-the-art technology designed to replicate the structure and function of specific organs or tissues in the human body on a miniature scale [50]. This technology offers a means of simulating the behavior of organs or tissues in a controlled and highly customizable environment [51]. One such organ model is the liver spheroid, a three-dimensional (3D) cellular structure or cluster of liver cells that closely mimics the architecture and function of the liver [39].

In our study, we employ the MPS system featuring liver spheroids to conduct biological experiments, leading to the generation of labeled spheroid images. This system is also integral to our liver toxicity assessment, as outlined in Section 3.3, where we expose the liver spheroids to different drug concentrations and observe their responses.

All liver spheroids used in this study were 3D InSight™ Human Liver Microtissues (InSphero AG), which comprised primary human hepatocytes and Non-Parenchymal Cells (NPC). The NPCs consisted of primary human Liver Endothelial Cells and Kupffer cells, obtained from multiple donors. The spheroids were cultured using the accompanying medium, 3D InSight™ Human Liver Microtissue Maintenance Medium – TOX (InSphero AG). Upon the spheroids’ arrival, the subsequent stages of our experiments were executed using the Company MPS platform.

To quantify spheroid viability, we employed the CellTiter-Glo^®^ 3D Cell Viability Assay (Promega, Cat# G9682), following the manufacturer’s protocol. After removing the culture medium from the Akura™ 96-well plates, each well was incubated with 50 µL of a 1:1 mixture of CellTiter-Glo 3D reagent and PBS for 30 minutes at room temperature on an orbital shaker. The lysates were then transferred to a Corning 96-well flat-bottom plate (Cat# 3693), which also contained a series of wells with known ATP concentrations used to construct a calibration curve. Luminescence was measured using a Synergy H1 plate reader, and ATP concentrations (in nanomolar) were computed by interpolating luminescence values against the calibration curve. These total ATP values per spheroid served as direct viability indicators and were later normalized by the median ATP value of control spheroids within each study to mitigate batch effects.

#### Study Description

To capture a diverse range of viability levels, multiple studies were conducted in which spheroids were allocated to specific treatment groups. These groups included both treatment and control spheroids. The treatment groups were subjected to a unique concentration of a drug, categorized as either toxic or non-toxic. The control spheroids did not receive any drugs; they were either in the same percentage of Dimethyl Sulfoxide (DMSO) as the treatment group plus medium, or solely in medium. Following a 1- to 7-day incubation period, which allowed for drug action in the treatment groups and stability in the control groups, the spheroids were imaged. Their ATP concentrations were quantified using ‘Promega CellTiter-Glo 2.0’[5], adhering to the manufacturer’s guidelines. This procedure yielded paired data of images and corresponding ATP values.

Figure 9 illustrates the study schedule, including spheroid acclimation on day 0, followed by various dosing, media exchange events, and terminal bioluminescence viability assays. The viability assay results, after control-normalization, are used as labels for training the viability regressor.

**Figure 9:**
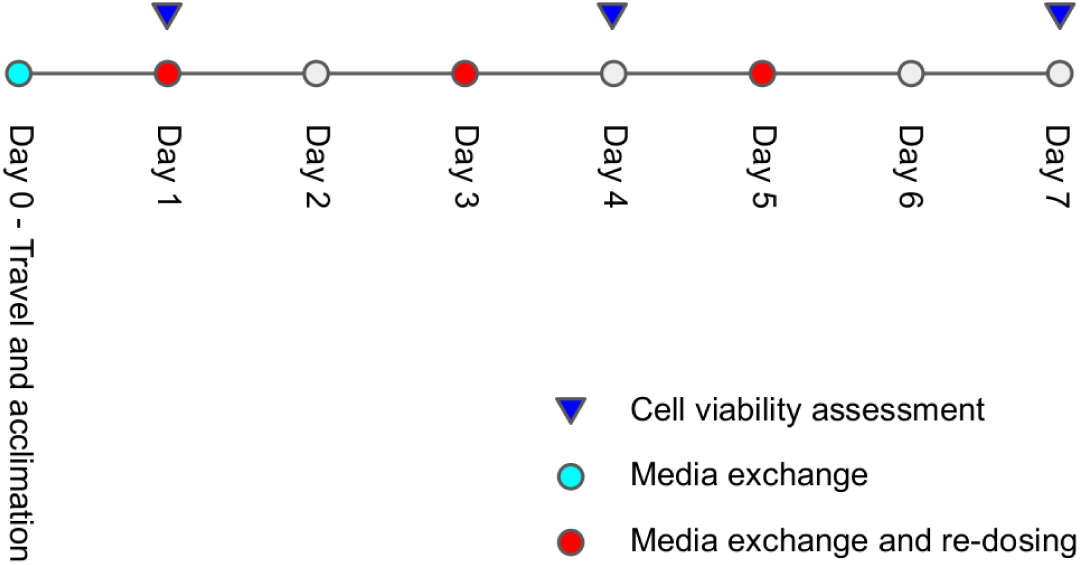
Study schedule detailing dosages and viability assessment regimens.

#### Dataset Description

The dataset comprises 2,359 labeled and 17,449 unlabeled images and is made available for further community research. Table 3 describes the four domains spanned by the dataset and their purposes for this study. Mainly, we use domain *A* as a train/validation/test domain, and the rest as purely test domains. An additional 17,449 unlabeled images are used for the DILI prediction study in Section 3.3. Each sample in the dataset represents an image together with its corresponding ATP value as quantified by our platform. The dataset is accompanied by a CSV file, the columns of which are described in Table 4.

**Table 3:** Breakdown of the viability dataset by domains.

| domain | laboratory | microscope | # images | split |
| --- | --- | --- | --- | --- |
| A | lab A | Echo Rebel | 1022 | train/val/test |
| B | lab B | Echo Revolve | 712 | test |
| C | lab A | Cytation | 245 | test |
| D | lab A | Thorlabs | 380 | test |

**Table 4:** CSV columns describing dataset samples.

| column | type | description | value examples |
| --- | --- | --- | --- |
| domain | str | Table 3 | A/B/C/D |
| location | str | laboratory id | A/B |
| study | str | study id | MS11 |
| day | int | day of imaging | 0..7 |
| treatment | str |  | drug/control |
| compound | str | compound administered | tolcapone |
| concentration_μM | float | conc. of compound | 0.3 |
| plate | str | plate id | 1197 |
| well | str | well id inside Akura96 plate | A6 |
| microscope | str | microscope model | echo_rebel |
| location_in_bucket | str | path to image | ./img.png |
| value | float | Measure viability in nM | 801.3 |
| w | int | image width | 2400 |
| h | int | image height | 2400 |
| xmin,ymin,xmax,ymax | float | 4 columns, spheroid position | 0.51, 0.42, 0.72, 0.61 |
| cytoplasmic_score | int | pathologist score | 0 |
| nuclear_score | int | pathologist score | 2 |
| contour_score | int | pathologist score | 4 |

The dataset is organized into multiple studies, each spanning a duration of 7 days. Lab A conducted 13 studies (7 labeled with viability measurements, 6 unlabeled), while Lab B conducted 5 studies. Within each study, the spheroids were divided into drug treatment groups and control groups. The drug treatment groups received specific doses of a drug along with DMSO used as a vehicle, whereas the control groups consisted of spheroids cultured in a pure medium or a medium supplemented with DMSO, but without any further treatments.

### 5.2 Batch Effect

The dataset analysis revealed the presence of batch effects, both across and within different studies. For example, Figure 10 illustrates the ATP values of control spheroids in different studies. Evidently, there is high variability in the ATP distribution of control spheroids on day 1 of the experiments.

**Figure 10:**
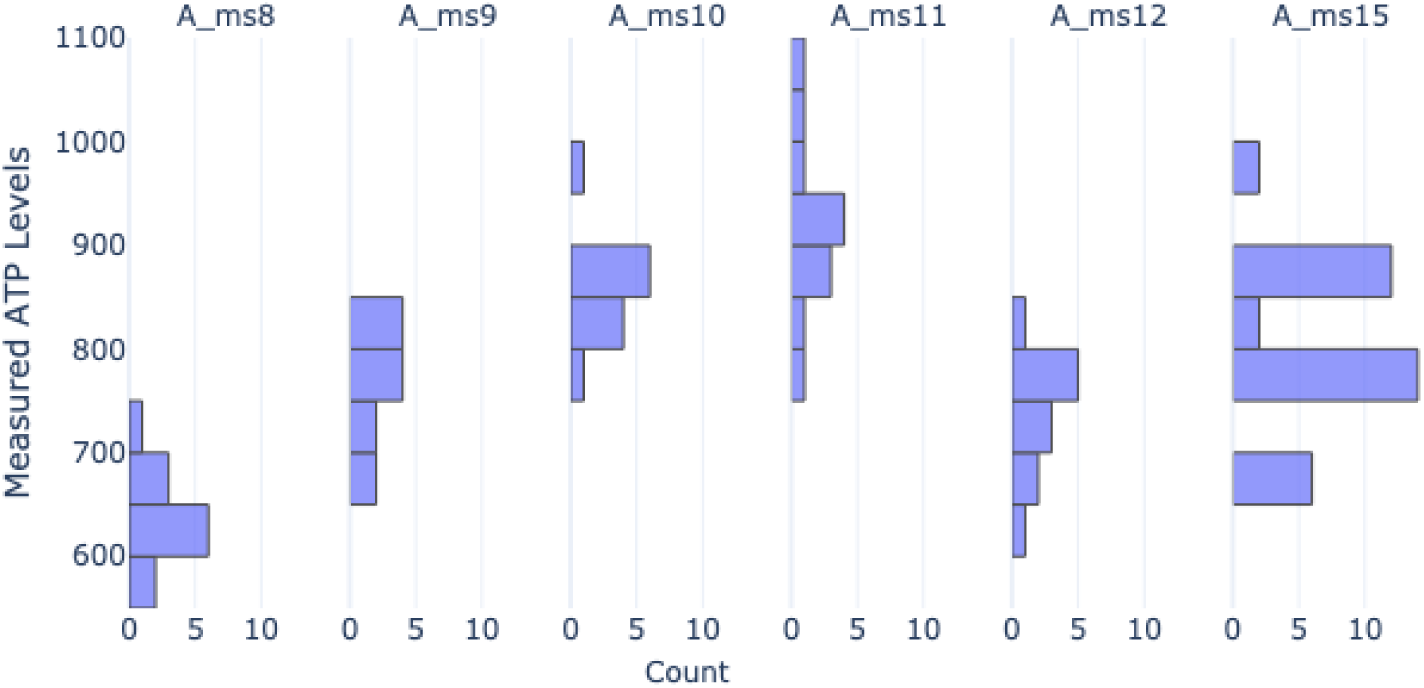
Observed batch effect between studies: Histograms of absolute viability values on day 1 for control spheroids in different studies.

Within individual studies, batch effects were observed based on the grouping of spheroids during the fluorescent sensing process. Variations in imaging data and viability assay results were found to be influenced by factors such as the timing of plate reading after the addition of the ATP-binding reagent during the assay protocol. These variations could result in different ATP measurements due to the reagent binding to varying numbers of ATP molecules over time. Additionally, other sources of batch effects within studies include human error, environmental factors, and potential variations in spheroid viability within different batches:

- Human error: Inaccurate pipetting during the addition of drug treatments or the execution of the viability assay protocol can lead to different amounts of medium or drug being introduced, affecting ATP measurements.
- Environmental factors: Factors such as temperature variations in the laboratory and fluctuations in lighting conditions during imaging can contribute to batch effects by impacting image quality and viability assay results.
- Spheroid starting conditions: Spheroids arriving in different batches may exhibit variations in viability and size.
- Laboratory equipment: Different microscopes or configurations can result in variations during data acquisition.

### 5.3 ATP Normalization

A key component of our method is a normalization preprocessing step, which addresses distribution shifts caused by batch effects and aligns the viability distributions more consistently. This alignment is visually demonstrated in Figure 11, where the distribution of viability from four independent studies is depicted before and after the normalization process. Specifically, the figure illustrates how the absolute viability samples, when normalized using their respective control counterparts, result in aligned distributions post-normalization, in contrast to the varied distributions observed pre-normalization. Given data sample *i*, which is a pair (*X_i_*, *y_i_*) in which *X_i_* is an image of a spheroid and *y_i_* is the measured ATP, we fetch the ATP values of all relevant control samples (*y*_1*c*_, *y*_2*c*_, …, *y_nc_*). The control-normalized value of 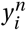 is then given by:

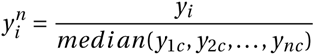

**Figure 11:**
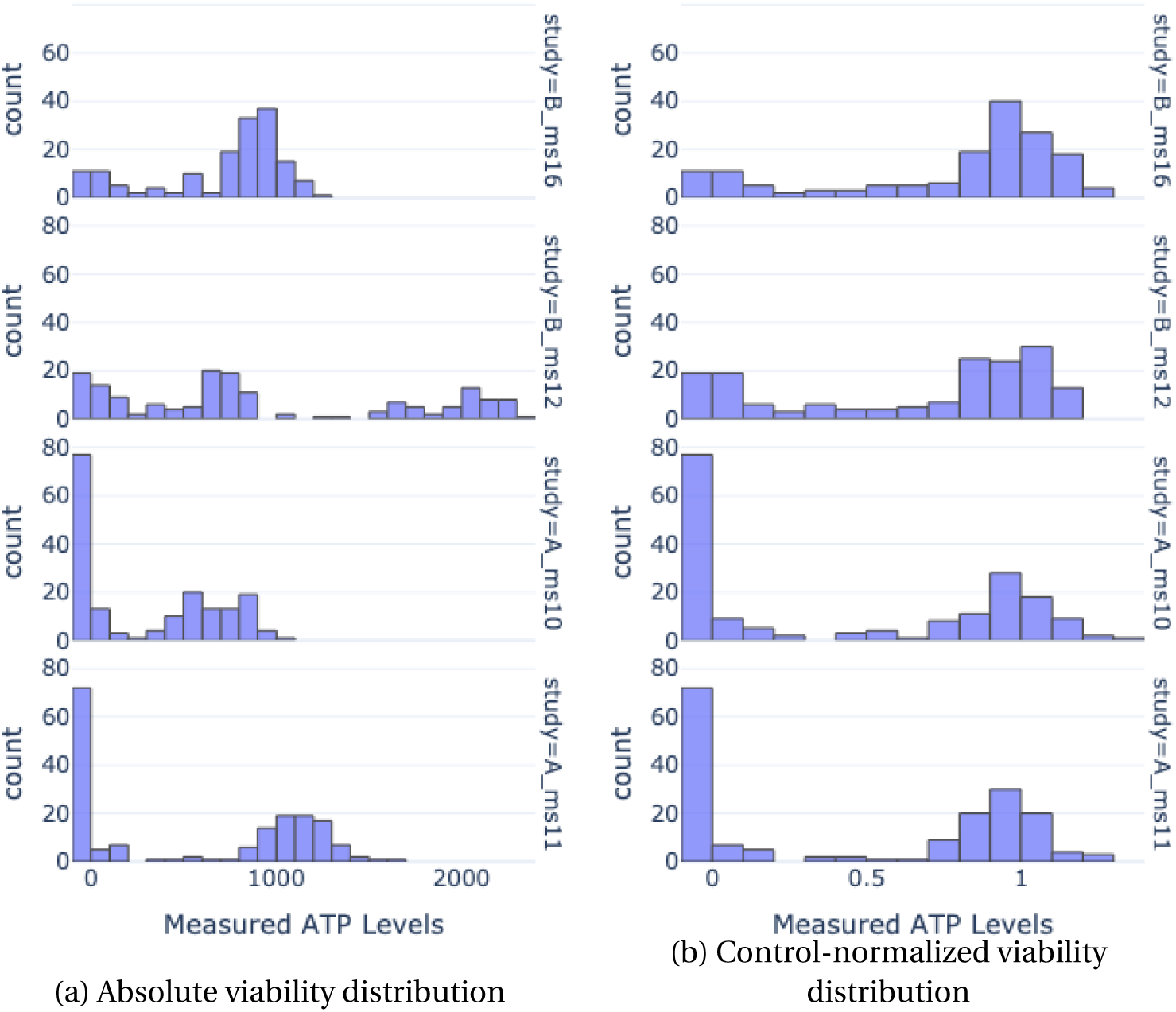
Distribution of viability from 4 independent studies before (panel (a)) and after (b) normalizing absolute viability samples by their respective control counterparts. Post-normalization, the viability distributions are aligned, as opposed to prenormalization.

This normalization by the median of control ATP values mitigates the influence of outliers in the control data, leading to more robust and reliable assessments. The effectiveness of this normalization is evident in Figure 11, as it visually represents the standardization achieved across different studies, thereby underscoring the importance and impact of this preprocessing step.

### 5.4 Architecture

#### NViR

We use a ResNet18[28] backbone, pre-trained on ImageNet[52], followed by an MLP regression head with 4 hidden layers of size 32. Overall, the model has 11 million learnable parameters. The backbone utilizes batch normalization[53] layers and ReLU activations. The regression head is regularized during training with 30% dropout [54]. Table 5 compares the performance of several different architectures used as backbones. All code is available at https://github.com/DanielDubinsky/atp_paper

**Table 5:**
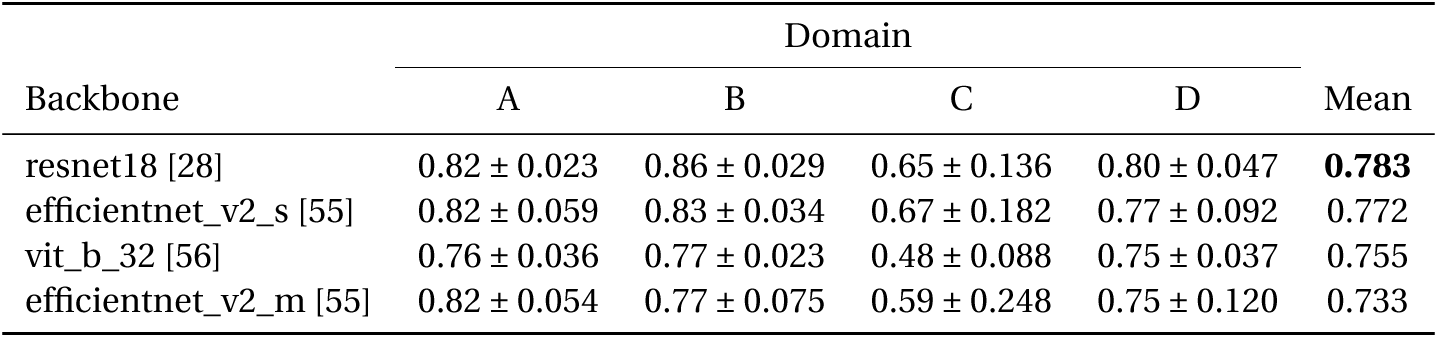
Mean and Standard Deviation for *R*^2^ (higher is better) of NViR with different backbones across four domains, with all models trained exclusively on domain *A*.

#### Baselines

All baselines are constructed in the same manner. For each baseline, 5-fold cross-validation on the training set is used to determine the best model type among DecisionTreeRegressor, GradientBoostingRegressor, and MLPRegressor. Various options for hyperparameters are explored for each model type to ensure optimal performance. For all baselines, GradientBoostingRegressor provided the best results in cross-validation, which is consistent with the literature on tabular data classification [57]. The GradientBoostingRegressor is a machine learning algorithm that constructs a predictive model in the form of an ensemble of weak prediction models, typically decision trees. It operates by building trees in a sequential manner, where each new tree aims to correct the errors made by the previously trained trees. The hyperparameters selected for each baseline are as follows:

- CVE baseline - 0.1 learning rate, 2 max depth, 4 min samples leaf, 2 min samples split, 50 n estimators.
- Pathology baseline - 0.1 learning rate, 3 max depth, 2 min samples leaf, 6 min samples split, 50 n estimators.
- CVE + pathology baseline - 0.1 learning rate, 2 max depth, 1 min samples leaf, 6 min samples split, 50 n estimators.

### 5.5 Training

The training architecture of NViR is depicted in Figure 12. During training, for each sample, all control spheroids are fetched, and the median of their ATP values is then used to normalize the sample’s ATP value, which is treated as the label.

**Figure 12:**
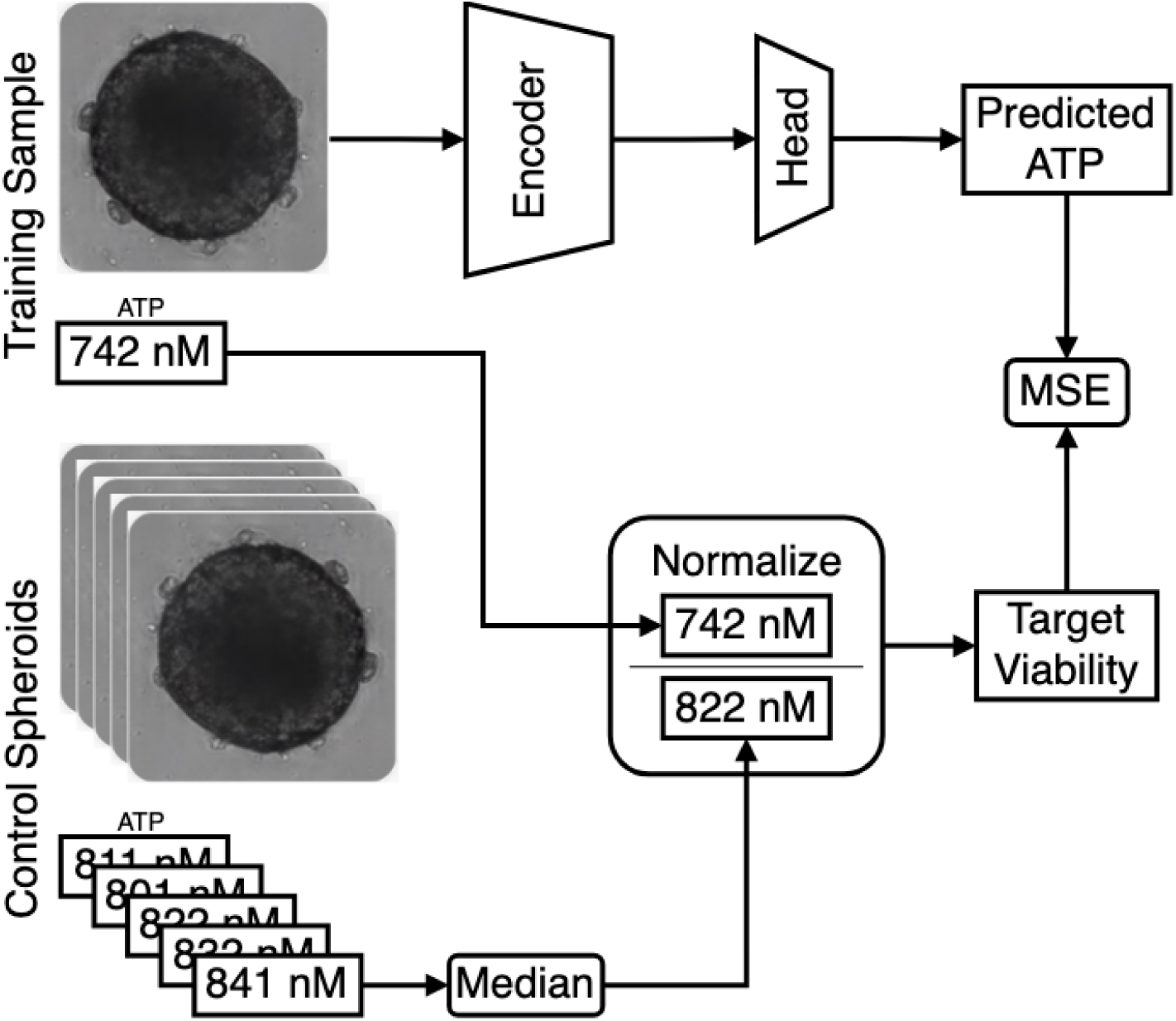
NViR training architecture.

All models were trained only on domain A. Domain A was split into training/-validation and test sets chronologically: all studies conducted before 2022-07-28 are considered for training/validation, while those conducted after are considered for testing. This split resulted in 777 training/validation images and 245 test images. Additionally, the test set included 712 images from domain B, 245 images from domain C, and 380 images from domain D. All ATP values are control-normalized, as described in Section 5.3.

#### Training

The optimized loss term is Mean Squared Error (MSE), with a batch size of 32, Adam optimizer[58], a learning rate of 0.0001 with a learning rate scheduler that reduces the rate by half after a plateau of 5 epochs, and a weight decay of 0.0005. Early stopping is employed, monitoring the validation loss with a patience of 20 epochs, and a maximum of 150 epochs. Training takes approximately 30 minutes on a single Tesla T4 GPU. Nine models were trained with three different initialization seeds and three different train/validation data split seeds - 80% for training and 20% for validation. The variation presented in Tab. 1 and 2 arises from applying the nine different models.

### 5.6 Classic Vision Encoder Features

In order to compute the classical computer vision features, we employ a binary segmentation mask that delineates the spheroids. These are obtained from a U-Net model [59] that we trained on a manually labeled subset of 1000 images. This model is relatively accurate, with an IoU above 0.9 over a validation set.

Binary masks of the spheroids, along with the images, are used to calculate the following visual properties of spheroids. These properties were chosen based on a description of how a domain expert, such as a pathologist, analyzes the morphology of spheroids in the context of viability estimation:

#### 1. Curvature

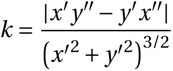

where *γ*(*t*) *=* (*x*(*t*), *y* (*t*)) is a parametric representation of the contour of the spheroid, as returned by cv2.findContours() on the mask, and *x′* and *y ′* are the gradients along the x and y axes, respectively. Intuitively, curvature describes how much the direction of a curve changes. Since healthy spheroids are smooth and resemble a circle when imaged, curvature is a property that can capture morphological changes where spheroids disintegrate and deviate from their original circular shape.

#### 2. Circumference / Perimeter

The ratio between the circumference of a circle with the same area as the spheroid and the perimeter of the spheroid. This ratio quantifies the closeness of the spheroid’s shape to a circle.

#### 3. Brightness

The ratio between the brightness of the spheroid and the background brightness, where brightness is the mean value of the pixels. As described by the pathologist, phenomena such as apoptosis are correlated with the darkening of the spheroid in bright-field images.

### 5.7 Pathologist Scoring Method

A Board Certified toxicological pathologist (AN) evaluated the bright-field images. The histopathological scoring was conducted in a blinded manner (i.e., the pathologist was not aware of the treatment or measured ATP values). The semi-quantitative scoring of five grades (0-4) [60] reflects the predominant degree of the specific lesion observed in the entire field of the histology section. For ease of interpretation, the scores were inverted in the analysis. The parameters scored were as follows:

- Cytoplasmic changes (i.e., presence of pallor, suggested to reflect the amount of glycogen). A higher degree (score) of cytoplasmic pallor is suggested to reflect a lower degree of stress effect. Conversely, a lower degree (score) of cytoplasmic pallor is suggested to reflect a high degree of stress effect.
- Nuclear changes (i.e., presence of apoptosis - pyknosis, karyorrhexis, or apoptotic bodies).
- Organoid contour shape changes, ranging from Grade 0, indicating no organoid changes (i.e., round-spherical), to Grade 4/marked, reflecting an indented contour shape.

Any degree of nuclear apoptotic changes, as well as changes in the organoid contour shape, are suggested to reflect reduced vitality (necrosis) and therefore should be considered adverse, according to the criteria of the American and European Societies of Toxicologic Pathology [61, 62]. Also, lower degrees (scores 0 to 2) of cytoplasmic pallor (glycogen) are judged to reflect a higher degree of stress and are therefore considered as reflecting adversity.

Figure 13 depicts the annotation platform used by the pathologist. The platform presents two images at each turn: the control spheroid (right) and the query spheroid (left). The pathologist then scores the query spheroid by each of the three parameters into one of the five levels. The pathologist-labeled dataset is published in [63].

**Figure 13:**
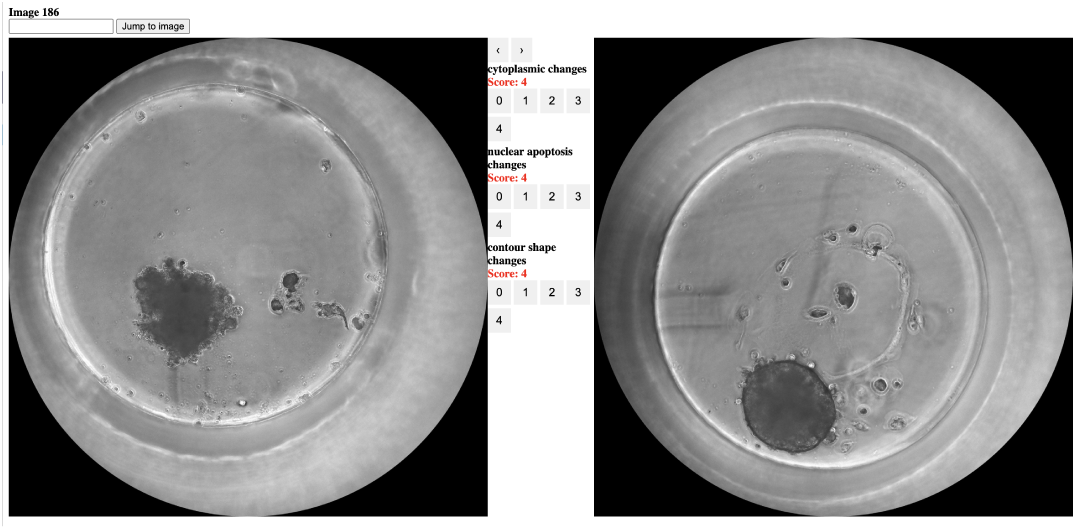
Screenshot of the annotation tool presented to the pathologist. The right image is the control spheroid, the left image is the query spheroid, and the interface between the images is for the actual annotation.

## 7 Data Availability

Data for this article are available at [63] at https://doi.org/10.5281/zenodo.15120118 Code is available at https://github.com/DanielDubinsky/atp_paper

## 8 Author Contributions

D.D. constructed and curated the machine learning dataset, conducted the experiments and statistical analysis, wrote the code, and drafted the manuscript. S.H. assisted in drafting and critically reviewing the manuscript. A.B. oversaw and provided guidance for the biological studies. S.Y. performed the biological experiments. B.K. and F.A. performed the biological experiments and contributed to the critical review. I.B. contributed to the critical review. A.N. contributed as a pathology domain expert to data annotation and critically reviewed the manuscript. L.W. supervised the work and contributed to both drafting and critically reviewing the manuscript.

## Acknowledgements

We would like to thank QurisAI, Y .Haran, the data science team (R. Bernstein, N. Schweiger, Y. Geffen) for valuable advice, the biology team (D. Zlotnik, A. Alon, G. Tenzer, L. Farberov, D. Rezania) and technicians (G. Margalit, D. Dubok,S. Jacobs,Y. Yohananov) for performing and monitoring the biological experiments and the engineering team (T. Abutbul, Y. Lowenhardt, O. Peleg, R. Divon, S. Gal, D. Haba) for setting up the MPS platform.

